# Approaching Maximum Resolution in Structured Illumination Microscopy via Accurate Noise Modeling

**DOI:** 10.1101/2023.12.07.570701

**Authors:** Ayush Saurabh, Peter T. Brown, J. Shepard Bryan, Zachary R. Fox, Rory Kruithoff, Cristopher Thompson, Comert Kural, Douglas P. Shepherd, Steve Pressé

## Abstract

Biological images captured by microscopes are characterized by heterogeneous signal-to-noise ratios (SNRs) due to spatially varying photon emission across the field of view convoluted with camera noise. State-of-the-art unsupervised structured illumination microscopy (SIM) reconstruction algorithms, commonly implemented in the Fourier domain, do not accurately model this noise and suffer from high-frequency artifacts, user-dependent choices of smoothness constraints making assumptions on biological features, and unphysical negative values in the recovered fluorescence intensity map. On the other hand, supervised methods rely on large datasets for training, and often require retraining for new sample structures. Consequently, achieving high contrast near the maximum theoretical resolution in an unsupervised, physically principled, manner remains an open problem. Here, we propose Bayesian-SIM (B-SIM), an unsupervised Bayesian framework to quantitatively reconstruct SIM data, rectifying these shortcomings by accurately incorporating known noise sources in the spatial domain. To accelerate the reconstruction process, we use the finite extent of the point-spread-function to devise a parallelized Monte Carlo strategy involving chunking and restitching of the inferred fluorescence intensity. We benchmark our framework on both simulated and experimental images, and demonstrate improved contrast permitting feature recovery at up to 25% shorter length scales over state-of-the-art methods at both high- and low-SNR. B-SIM enables unsupervised, quantitative, physically accurate reconstruction without the need for labeled training data, democratizing high-quality SIM reconstruction and expands the capabilities of live-cell SIM to lower SNR, potentially revealing biological features in previously inaccessible regimes.

## 1 Introduction

Fluorescence microscopy is a method of choice for studying biological processes in live cells with molecular specificity. However, the diffraction of light limits the experimentally achievable resolution for conventional microscopy to ≈200 nm [1], setting a lower bound on the microscope’s ability to probe the molecular events underlying life’s processes. In response, there is strong interest in developing experimental and computational superresolution methods that extend microscopy beyond the diffraction limit. Unfortunately, to date, most superresolution techniques strongly constrain the sample geometry, sample preparation, or data collection strategy, rendering superresolved live-cell imaging challenging or impossible. For example, stimulated emission depletion microscopy (STED) [2] requires high laser power and point-scanning, although recent parallelized approaches have expanded its capabilities [3]. Localization- [4, 5, 6, 7] and fluctuation-based [8, 9] techniques require specific fluorophore photophysics and hundreds to thousands of images, limiting their applicability to live-cell imaging. Structured illumination microscopy (SIM) [10, 11] is an attractive alternative for superresolved live-cell imaging because it is compatible with standard sample preparation methods, requires only about 10 raw images per frame, and uses lower illumination intensity leading to reduced phototoxicity and photobleaching.

In SIM, structured illumination of a sample, combined with incoherent imaging, translates high-frequency information from beyond the diffraction limit below the microscope band-pass, where it appears as moiré interference fringes. In linear SIM, this additional information is used in computationally reconstructing the underlying fluorescence intensity map with theoretically up to double the widefield resolution. By exploiting photophysics, non-linear SIM can achieve even larger resolution enhancements [12, 13]. Many 2D SIM approaches use a set of nine sinusoidal illumination patterns, both rotated and phase-shifted, to generate near isotropic lateral resolution improvement. However, other 2D patterns or random speckle patterns are also used for similar purposes [14, 15] often generated using gratings [16] or spatial light modulators [17, 18].

The computational reconstruction framework used in recovering information beyond the microscope’s traditional spatial resolution limit often determines the effective resolution in SIM, due to the choices made in how to treat contrast degrading noise.

Approaching the maximum recoverable resolution in the SIM reconstruction as warranted by the data, therefore, requires correctly treating the main sources of noise, Poissonian photon emission and camera electronics [19, 20] by accurately modeling the physics of image formation.

Furthermore, the inherently stochastic nature of fluorescence imaging makes reconstructing the fluorescence intensity map, the product of the fluorophore density and the quantum yield, naturally ill-posed. Consequently, we must use a model that provides statistical estimates of the fluorescence intensity map from exposure-to-exposure fluctuations in photon counts. In the high-SNR regime, where photon shot noise dominates camera noise, average photon counts allow accurate estimation of the noise statistics from the recorded pixel counts. The ill-posedness of a reconstruction is, therefore, less severe at high-SNR.

However, fluorescence microscopy images often include low-SNR regions due to low fluorophore density or low illumination power. Image-to-image fluctuations are large in such regions, and both photon shot noise and camera noise become important, making it more challenging to estimate the fluorescence intensity map. For incoherent imaging methods such as SIM, the effective SNR also depends on the length scales of features present in the image. The SNR-spatial size relationship is naturally understood in the Fourier domain where short length scales correspond to high spatial frequencies. As a microscope’s incoherent optical transfer function (OTF) attenuates high spatial frequency information [21], there is loss of contrast for small spatial features, regardless of the average photon count. The inherent loss of contrast worsens the ill-posedness of image reconstruction and makes it very challenging to distinguish small biological features from pixel-to-pixel variations in photon shot and camera noise. Put differently, the effective SNR in the recorded images decreases with increasing spatial frequency, making the high spatial frequency information especially susceptible to noise.

Image reconstruction challenges are further compounded in SIM where super-resolved fluorescence intensity maps are recovered by unmixing and appropriately recombining noisy, low-contrast information distributed through the collected raw images as shifting moiré fringes. However, image-to-image variation due to photon shot and camera noise at the single pixel level can produce intensity fluctuations indistinguishable from low-contrast fringes. Consequently, during the SIM reconstruction process, these noisy signals may masquerade as genuine moiré fringes, leading to high-frequency reconstruction artifacts such as hammerstroke [22]. Artifacts typically become particularly pronounced at low-SNR and for moiré fringes near the diffraction cutoff frequency, where the OTF reduces the contrast as discussed earlier.

Multiple SIM reconstruction tools have been developed to limit artifacts in SIM reconstructions. The most common reconstruction algorithms [23] rely on a Wiener filter to deconvolve and recombine raw images in the Fourier domain. However, these approaches may fail in low-SNR regions and can introduce artifacts indistinguishable from real structures [24, 25] due to their sub-optimal treatment of noise. For instance, performing a true Wiener filter requires exact knowledge of the SNR as a function of spatial frequency. Unfortunately, it is difficult to estimate the exact relationship of SNR versus spatial frequency from the sample. Instead, approaches replace the true SNR with the ratio of the magnitude of the OTF and a tuneable Wiener parameter [26]. More sophisticated approaches attempt to estimate the SNR by assuming the sample signal strength obeys a power-law [27], choosing frequency-dependent Wiener parameters based on noise propagation models [28], or alternatively engineering the OTF to reduce common artifacts [29].

The primary difficulty associated with these approaches, and what ultimately limits feature recovery near the maximum supported spatial frequency, is that the noise model is well understood in the spatial domain while SIM reconstructions are performed in the Fourier domain. Rigorously translating the noise model to the Fourier domain is challenging because local noise in the spatial domain becomes non-local in the Fourier domain and gets distributed across all spatial frequencies.

Many attempts at avoiding artifacts associated with Wiener filter based approaches at low-SNR [23] have proven powerful, yet they still treat noise heuristically and typically require expensive retraining or parameter tuning to reconstruct different biological structures or classes of samples such as mitochondrial networks [30], nuclear pore complexes [31], and ribosomes [32]. For example, one common strategy is to apply content-aware approaches relying on regularization to mitigate ill-posedness of the reconstruction problem by *e*.*g*., smoothing the fluorescence intensity map, or using prior knowledge of the continuity or sparsity of the biological structures. These approaches include TV-SIM [33], MAP-SIM [34], Hessian-SIM [35, 36], and proximal gradient techniques [37, 38, 39]. Another recently developed self-supervised strategy uses implicit image priors such as deep image priors (DIP) [40, 41, 42, 43] and physics informed neural networks (PINN) [44] that are neural networks whose structure inherently constrains the output images. These networks are typically optimized iteratively using the input raw images only but can have a tendency to overfit data with additional regularizers. On the other hand, recently developed deep-learning based 2D-SIM approaches including rDL-SIM [45] and DFCAN/DFGAN [46] train neural networks in a supervised fashion to recognize structures at the cost of requiring large datasets for training and have been demonstrated only for TIRF-SIM modalities, to avoid out-of-focus background. While these methods are powerful, developing deep learning approaches that generalize to different SIM instruments and classes of samples remains an open challenge. Alternative approaches are therefore greatly needed that operate at low-SNR and thus broadly apply to any mixed-SNR sample type.

Here we present Bayesian structured illumination microscopy (B-SIM), a new SIM reconstruction approach capable of resolving features near the resolution limit warranted by data, and handling high-and low-SNR SIM data in a fully principled way without the need for assumptions about biological structures or labeled training data. Our proposed approach addresses many of the deficiencies of previous SIM algorithms discussed above and offers a powerful general purpose alternative to deep learning approaches. Specifically, B-SIM: 1) accurately models the stochastic nature of image formation process; 2) permits only positive fluorescence intensities; 3) eliminates arbitrary constraints on the smoothness of biological features; and 4) is amenable to parallelized computation. These features are achieved by working in a Bayesian paradigm where every part of the image formation process, including Poissonian photon emission and camera noise, are naturally incorporated in a probabilistic manner and used to rigorously compute spatially heterogeneous uncertainty.

This advancement in noise modeling results in contrast enhancement near the highest supported spatial frequencies permitting feature recovery in low-SNR data at up to 25% shorter length scales than state-of-the-art unsupervised methods. Previous Bayesian approaches [47] incorporated stochasticity but until now have not correctly incorporated physics due to their choice of Gaussian process priors that enforce spatial correlations on biological features, permit negative fluorescence intensity values, and do not apply at low-SNR where the Gaussian approximation to the full noise model falters.

## 2 Results

B-SIM operates within the Bayesian paradigm where the main object of interest is the probability distribution over all fluorescence intensity maps ***ρ***, termed posterior, as warranted by the data. The posterior is constructed from both the likelihood of the collected raw SIM images given the proposed fluorescence intensity map together with any known prior information such as the domain over which the fluorescence intensity map is defined via a prior probability distribution (see methods). From the posterior probability distribution, we compute the mean fluorescence intensity map ⟨***ρ***⟩ that best represents the biological sample. Furthermore, the posterior naturally provides uncertainty estimates for the fluorescence intensity maps, reflecting spatial variations in noise due to any heterogeneity.

As shown in Fig. 1, to learn the posterior over the fluorescence intensity maps, ***ρ***, we first develop a fully stochastic image formation model taking into account photon shot noise and CMOS camera noise. This model allows us to formulate the probability of a candidate fluorescence intensity map given input raw images, a calibrated point-spread-function (PSF), illumination patterns, and camera calibration maps. As our posterior does not attain an analytically tractable form, we subsequently employ a requisite Markov Chain Monte Carlo (MCMC) scheme to draw samples from the posterior, and compute its mean and associated uncertainty mentioned earlier. Furthermore, we parallelize this sampling scheme by first noting that a fluorophore only affects a raw image in a small neighborhood surrounding itself owing to the PSF’s finite width. By dividing images into chunks and computing posterior probabilities locally, our parallelized approach avoids large, computationally expensive convolution integrals.

**Fig. 1:**
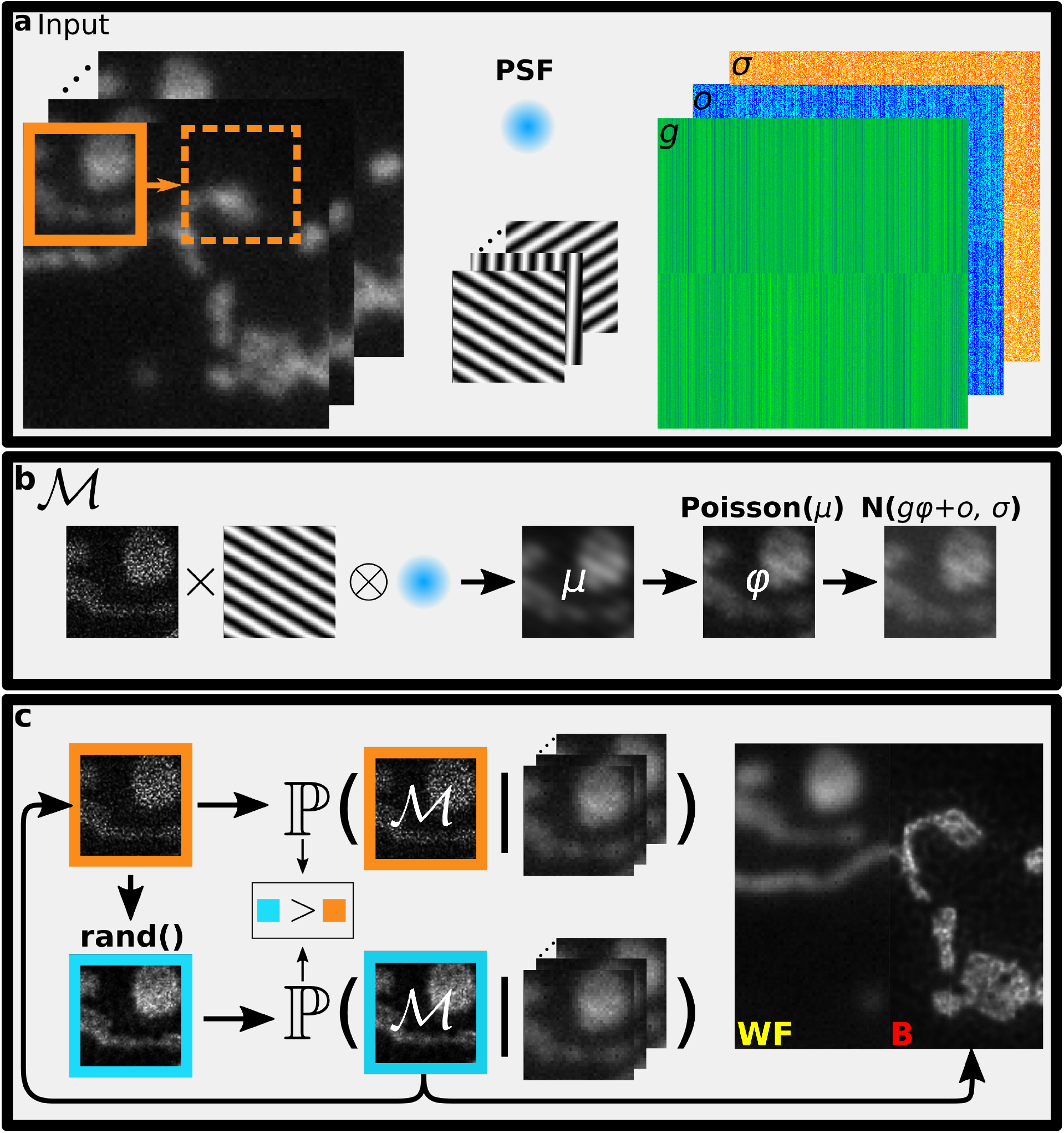
Bayesian-SIM reconstruction with spatial domain noise propagation. **a**. SIM involves collecting fluorescence images (left) illuminated by structured intensity patterns (center). We utilize a point-spread function model (center)(PSF) and calibrate the camera noise parameters (right): gain (g), offset (o), and readout noise (σ). To infer the underlying fluorescence intensity map, we start from a small region (orange) and sweep over the entire sample in a parallelized fashion. **b**. An image formation model, ℳ, involves multiplication of an underlying fluorescence intensity ma with illumination patterns and convolution with the PSF to obtain a noiseless image, µ, which is then corrupted by photon shot noise and camera noise. **c**. A Monte Carlo algorithm to sample candidate fluorescence intensity maps from the posterior probability distribution. Based on the current sample (orange) we propose a new sample for the fluorescence intensity map (blue) and compute its corresponding posterior probability. Favoring higher probability, we stochastically accept or reject the proposed fluorescence intensity map and update the current fluorescence intensity map. We average many samples of fluorescence intensity maps after convergence to obtain B-SIM reconstruction (right). WF and B denote widefield image and B-SIM reconstruction, respectively.

To validate B-SIM, we reconstruct SIM data in a variety of scenarios briefly highlighted here, and demonstrate that B-SIM surpasses the performance of existing algorithms both at low-SNR and, surprisingly, high-SNR. First, as proof of concept, we consider simulated fluorescent line pairs with variable spacing generated from a known imaging model and demonstrate the best performance achievable under ideal conditions. Next, we consider an experimental test sample with variable spaced line pairs. This demonstrates that B-SIM retains excellent performance under well-controlled experimental conditions. Finally, we analyze experimental images of fluorescently labeled mitochondria in live HeLa cells, where experimental imperfections in the illumination, out-of-focus background fluorescence, and pixel-to-pixel variation in camera noise introduce challenges in SIM reconstruction.

In each case, we compare B-SIM with reconstructions using Wiener-filter based methods, particularly HiFi-SIM [29], and optimization based approaches, including FISTA-SIM [48]. We find that B-SIM provides increased fidelity and contrast for short length scale features at both high- and low-SNR as compared to existing state-of-the-art approaches.

Extending beyond unsupervised approaches, we compared the performance of B-SIM to deep learning based rDL-SIM [45] pre-trained for publicly available TIRF-SIM microtubule data [49] provided by the authors of [46, 45] themselves and newly generated experimental TIRF-SIM microtubule data. Following the procedure from Ref. [45], we denoise the raw SIM images using the provided microtubule specific neural network. After denosing, we reconstruct SIM images using HiFi-SIM [29]. For both the BioSR and new experimental data, we find that B-SIM produces improved contrast as shown in supplementary Fig. S5.

### 2.1 Simulated Data: Line Pairs

To demonstrate B-SIM performance under well-controlled conditions, we simulated a fluorescence intensity map of variably-spaced line pairs with separations varying from 0 nm to 330 nm in increments of 30 nm. The sample is illuminated by nine sinusoidal SIM patterns with frequency 0.9 times that of Abbe diffraction limit, assuming NA = 1.49 and emission light wavelength λ = 500 nm. With these microscope parameters, the maximum supported spatial frequency for widefield imaging is 2 NA/λ ≈ (168 nm)^*−*1^, while the maximum recoverable frequency in SIM is 3.8 NA/λ ≈ (88 nm)^*−*1^. We generated simulated datasets at both high-SNR, where photon shot noise is the dominant noise source, and at low-SNR, where both shot noise and camera noise are significant.

For the high-SNR dataset, we generated raw SIM images with up to ≈1000 photons-per-pixel. In Fig. 2a-b, we show SIM reconstructions of the fluorescence intensity maps obtained using B-SIM and compare with HiFi-SIM, and FISTA-SIM. At high-SNR, all three approaches produce high-quality fluorescence intensity maps with minimal artifacts. We find that, according to the Sparrow criterion, the widefield image resolves the 180 nm line pair, HiFi SIM resolves the 120 nm line pair, and both FISTA-SIM and B-SIM resolve the 90 nm line pair, approaching the theoretical limit discussed above. To further assess the resolution and contrast enhancements obtained across various methods, we also plot the intensity along line cuts showing the line pairs separated by 90 nm and 120 nm. Here we clearly see that B-SIM produces significantly enhanced contrast for both line pairs, with over ≈40% intensity drop in the center for the 90 nm line pair as compared to ≈5% drop in the FISTA-SIM reconstruction. Indeed, in comparison with HiFi-SIM, B-SIM recovers significantly higher contrast for line pairs with separations <150 nm. We attribute the enhanced contrast obtained by B-SIM to its physically principled incorporation of noise as compared to other approaches.

**Fig. 2:**
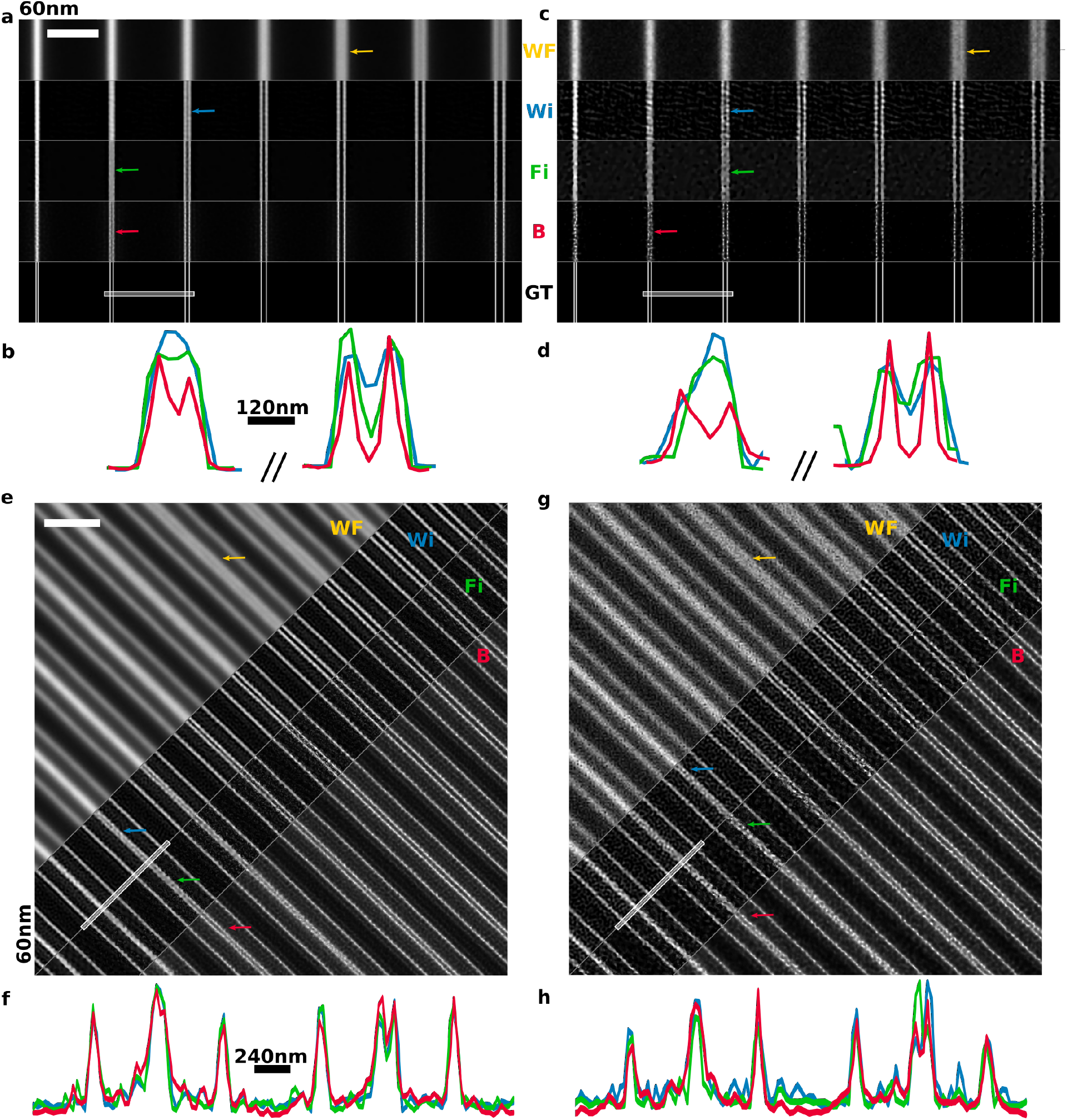
SIM reconstruction of variably spaced line pairs. **a**. Simulated line pairs with spacing ranging from 60 nm to 210 nm in steps of 30 nm at high-SNR. We show the pseudo-widefield (WF) image obtained by averaging raw data, and ground truth (GT) image, together with SIM reconstructions using Wiener (Wi), FISTA-SIM (Fi), and B-SIM (B). Scale bar 1.5 µm. Colored arrows indicate the line pair resolved according to the Sparrow criterion. **b**. Line cuts corresponding to white line in a. All reconstruction methods resolve the 120 nm-spaced line pair. Both FISTA-SIM and B-SIM resolve the 90 nm-spaced line pair, but B-SIM does so with higher contrast. **c**. Simulated line pairs as in a., but at low-SNR. **d**. Line cuts corresponding to c. All reconstruction methods resolve the 120 nm line pair, but only B-SIM resolves the 90 nm line pair. **e**. Experimental images and reconstructions of variably spaced line pairs on an Argo-SIM calibration sample at high-SNR. Line pairs have spacings of 60 nm to 330 nm in 30 nm steps. Scale bar 2.0 µm. **f**. Line cuts corresponding to e. All methods resolve the 120 nm-spaced line-pair (right) with similar contrast, and no methods resolve the 90 nm line pair (left). **g**. Experimental images as in e., but at low-SNR. Wiener and FISTA-SIM introduce reconstruction artifacts. **h**. Line cuts corresponding to h. Only B-SIM resolves the 120 nm spaced line pair.

We also tested B-SIM’s robustness against errors in illumination patterns for the high-SNR dataset. By adding erroneous shifts to the phases of the sinusoidal patterns used to compute the posterior, we find that the reconstruction is robust for phase errors up to order 10°. Significant artifacts appear in the reconstruction for larger errors, as shown in supplementary Fig. S1.

Next, we consider a more challenging low-SNR simulated dataset with up to ≈40 photons per pixel in each raw SIM image, Fig. 2c-d. At this signal level, photon noise results in larger exposure-to-exposure variance and camera noise becomes significant. Furthermore, a simple Poissonian or Gaussian approximation of the noise model is not enough for a high fidelity reconstruction. Due to the decreasing effective SNR with increasing spatial frequency, we expect the achievable contrast at shorter length scales to be lower than that of the high-SNR case. This intuition is confirmed in Fig. 2, where we find that wide-field only resolves the 210 nm line pair, instead of the 180 nm as in the high-SNR case. At low-SNR, SIM reconstruction becomes increasingly challenging as evidenced by increasing hammerstroke artifacts visible in the HiFi- and FISTA-SIM reconstructions. Here, both HiFi-SIM and FISTA-SIM are unable to resolve separations below 120 nm. However, we find B-SIM still resolves the 90 nm line pair without hammerstroke artifacts.

To quantitatively compare the various reconstruction methods, we also calculated the mean-square error (MSE) and peak signal-to-noise ratio (PSNR) using the known ground truth image, as shown in supplementary Table S1. For the high-SNR data, we find that both B-SIM and FISTA-SIM more faithfully reconstruct the line pairs than the other methods, and achieve similar scores. More critically, for the low-SNR data, B-SIM achieves the highest scores. To further improve the smoothness in the B-SIM results shown in Fig. 2a-d, it is possible to generate more MCMC samples, albeit at a higher computational cost.

### 2.2 Experimental Data: ArgoSIM Line Pairs

Having demonstrated significant contrast improvement by B-SIM on simulated data near the maximum achievable SIM resolution, we next recover fluorescence intensity maps from experimental images of variably-spaced line pairs on an ArgoSIM calibration slide. The line pairs are separated by distances varying from 0 nm to 390 nm in increments of 30 nm. We illuminated these line pairs with nine sinusoidal illumination patterns with three orientations, each with three associated phase offsets, and pattern frequency of ≈0.8 times that of Abbe diffraction limit. The peak emission wavelength for this sample was λ = 510 nm and objective lens had NA = 1.3. Here the diffraction limited resolution is ≈196 nm and achievable SIM resolution is ≈109 nm. With these parameters, we generated two datasets, one at high-SNR, and one at low-SNR per exposure.

Unlike in analyzing simulated line pairs where the illumination pattern is specified by hand, in working with Argo-SIM slides, we must estimate the illumination patterns from the raw fluorescence images as a pre-calibration step prior to learning the fluorescence intensity map. We estimate pattern frequencies, phases, modulation depths, and amplitudes from high-SNR images using Fourier domain methods [18], and generate one set of illumination patterns to be used for Wiener, FISTA-SIM, and B-SIM reconstructions. An independent approach for pattern estimation might not be feasible in all experimental circumstances. Joint pattern and fluorescence intensity map optimization is an interesting potential extension of B-SIM. We do not pursue this further in this work due to the increase in computational expense entailed in this joint inference.

For the high-SNR dataset, we generated raw SIM images with up to ≈200 photons-per-pixel, Fig. 2e-f. Here, all SIM reconstruction methods generate high quality fluorescence intensity maps with minimal artifacts. HiFi-, FISTA-, and Bayesian-SIM reconstructions all resolve the 120 nm line pair, with B-SIM resulting in the best contrast. Unlike for the simulated sample, none of the line pair spacings here are right at the diffraction limit, explaining why B-SIM does not resolve an extra line pair compared with other methods here.

For the low-SNR dataset, we generated raw SIM images with up to ≈40 photons-per-pixel by using a shorter illumination time. We found HiFi-SIM was not able to accurately estimate the illumination pattern parameters from this low-SNR data, so instead we use our mcSIM Wiener reconstruction [18]. As for the simulated data, Wiener-, and FISTA-SIM amplify high-frequency noise leading to significant reconstruction artifacts and these methods no longer clearly resolve the 120 nm line pair with high contrast, as shown in Fig. 2g-h. On the other hand, B-SIM introduces fewer high-frequency artifacts and continues to resolve the 120 nm line pair.

### 2.3 Experimental Data: Mitochondria Network in HeLa Cells

Next, we consider B-SIM’s performance on experimental images of dynamic mitochondrial networks in HeLa cells undergoing constant rearrangement in structure through fission and fusion with formations such as loops [50, 30, 51]. We perform live cell imaging to avoid mitochondria fixation artifacts [52] using the same experimental setup as for the ArgoSIM sample, but with 635 nm illumination and 660 nm emission peak. We also adjust the absolute SIM pattern frequency to remain fixed at ≈0.8 the diffraction limit. For this peak emission wavelength, the Abbe resolution is approximately ≈254 nm and maximum SIM resolution is ≈141 nm. As in the previous cases, we collect paired high- and low-SNR datasets.

In these datasets, due to high variation in fluorophore density along the mitochondrial network, the photon emission rate or SNR varies significantly throughout the image, posing a challenge for existing SIM reconstruction algorithms. Furthermore, while mitochondria are commonly imaged using fluorescent labels that emit near 500nm wavelength to gain resolution [53], the use of far-red fluorescent labels makes the data presented here particularly challenging for recovering small features in mitochondria networks. On the other hand, improvement in SIM reconstructions using such labels could be significantly advantageous because longer excitation wavelengths can reduce phototoxicity, background fluorescence, Rayleigh scattering, and Raman scattering [54].

For the high-SNR dataset, we generated raw SIM images with up to ≈150 photons-per-pixel, Fig. 3a-d. To clearly distinguish recovered superresolution information from artifacts, we image the mitochondrial network at two different time points 6.082 s apart, set by the total time required to collect multiple datasets with different illumination times and SNR (see methods). This time scale is fast compared with the network rearrangement and, as such, we do not expect major morphological changes to occur. Collecting paired images allows us to experimentally differentiate recovered superresolved information, which should appear at both time points, from reconstruction artifacts, which should not. For the high-SNR sample, HiFi- and FISTA-SIM produce reasonable reconstructions showing high-resolution features. FISTA-SIM introduces reasonably strong staircase artifacts [55]. On the other hand, B-SIM achieves significantly better contrast at short length scales, more clearly revealing mitochondrial morphology. To illustrate the differences in these reconstructions, we focus on a tubular loop approximately 195 nm in diameter (Fig. 3b,d) and display a line cut. This loop appears in the network in B-SIM reconstructions at both time points. HiFi-SIM, on the other hand, is unable to resolve the central dip in fluorescence within the loop.

**Fig. 3:**
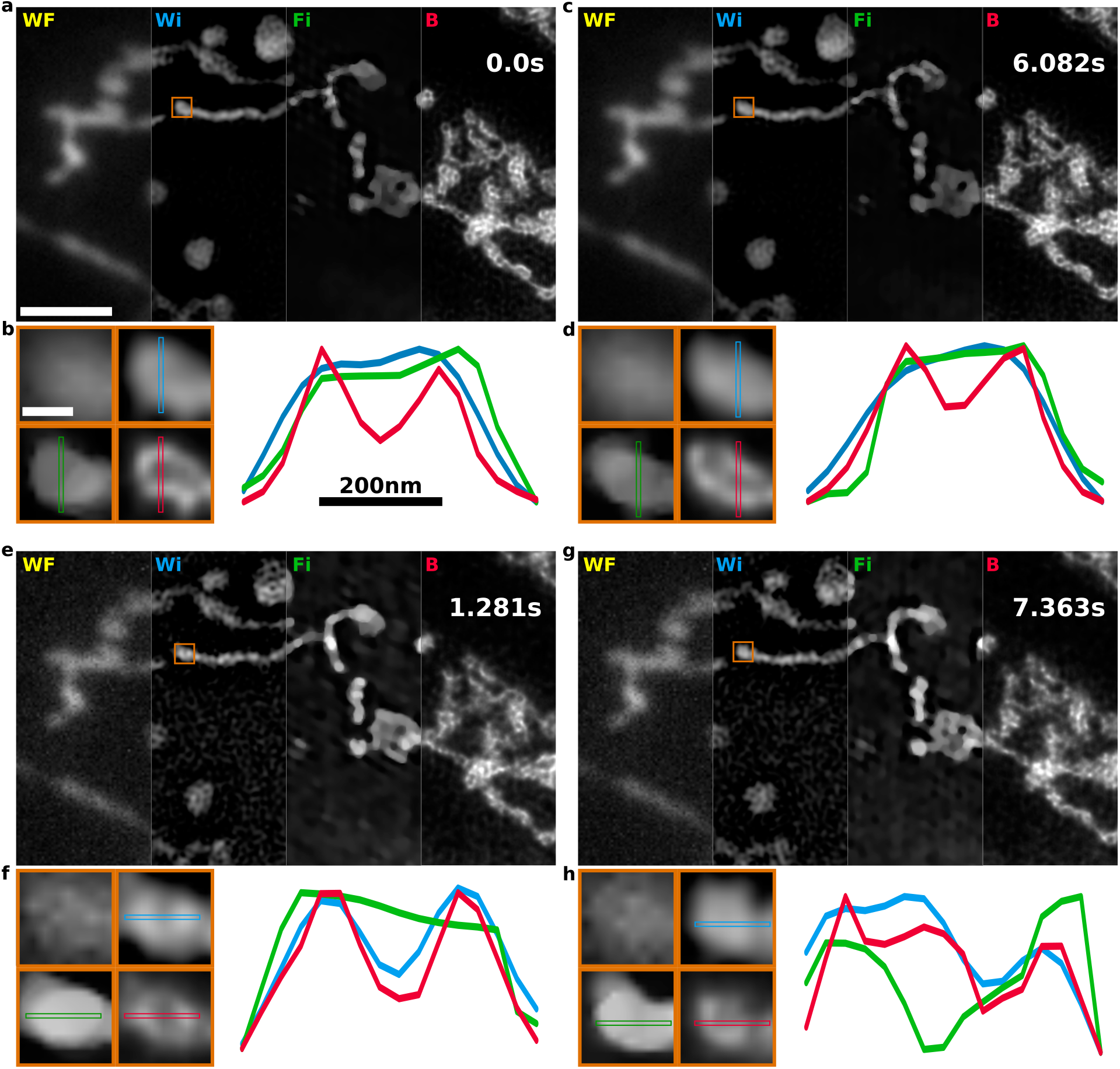
SIM reconstruction of mitochondrial networks. **a**. Pseudo-widefield (WF) and SIM reconstructions of MitoTracker deep red labelled mitochondria in HeLa cells at high-SNR. We compare Wiener filter (Wi), FISTA-SIM (Fi), and B-SIM (B) reconstruction methods. At high-SNR, all methods capture superresolution information with limited reconstruction artifacts. Scale bar 2.5 µm. **b**. Region of interest corresponding to the orange box in a. (left) shown in widefield, Wiener, FISTA-SIM, and B-SIM. Scale bar 300 nm. Line cuts (right) demonstrate B-SIM recovers more superresolution information than other methods. **c**. Mitochondrial network ≈6 s later. The superresolved structures appear in the same locations, suggesting these are real features and not artifacts. **d**. Same region of interest as in b. **e**. Low-SNR SIM reconstruction of the same sample using a shorter illumination time. This image is acquired 1.281 s after the image in a. **f**. Regions of interest at low-SNR. **g**. Low-SNR image ≈6 s later. **h**. Regions of interest at low-SNR and second time-point.

Unlike for the variably-spaced line pairs, there are no features in the mitochondrial network of known size we can use to directly infer the resolution achieved by the various SIM reconstruction methods. To obtain a quantitative estimate, we rely on image decorrelation analysis [56]. We estimate the achieved widefield resolution to be 325 nm compared with 180 nm for the Wiener reconstruction, and 131 nm for B-SIM, indicating ≈25% improvement which broadly agrees with our observations in Fig. 3a-d. Lastly, we do not report results for FISTA-SIM as the decorrelation analysis relies on phase correlations in Fourier domain that are strongly affected by the total variation (TV) regularization.

We note that decorrelation analysis provides a resolution estimate of 131 nm, smaller than the Abbe limit of 141 nm. However, this does not mean that B-SIM achieves resolutions better than the Abbe limit. Because decorrelation analysis relies on phase correlations present in the raw data in Fourier space, it likely indicates that B-SIM effectively smooths the reconstruction on spatial scales of the order of, and slightly smaller than, the diffraction limit. Such smoothing may be related to the approximate Gaussian OTF used in this reconstruction, which does not have a hard cut-off spatial frequency. On the other hand, decorrelation analysis relies on identifying the maxima in correlation-versus-resolution curves for a sequence of high- and low-pass filters. Based on the difference in the peak locations for the curves in this sequence, we estimate the uncertainty in the decorrelation analysis resolution to be ≈5 nm, placing the Abbe limit nearly within the uncertainty estimate.

Lastly, for the low-SNR dataset, we generated raw SIM images with up to ≈30 photons per pixel, Fig. 3e-h. These images were interleaved with the high-SNR images, and so we expect the morphology to remain the same. The HiFi-SIM reconstruction shows high-frequency noise amplification artifacts. These are avoided in the FISTA-SIM by applying a strong regularizer, which however significantly reduces the achievable resolution. On the other hand, B-SIM results in high-resolution reconstruction. We consider the same mitochondrial loop structure as before, and find that it is resolved in both HiFi- and B-SIM. Applying decorrelation analysis, we find similar results as before where the widefield resolution is 430 nm, compared with 176 nm for the Wiener reconstruction, and 131 nm for B-SIM.

### 2.4 Uncertainty in B-SIM Reconstruction and Statistical Estimators

Bayesian inference using MCMC sampling techniques to draw samples of candidate fluorescence intensity maps from a posterior naturally facilitates uncertainty quantification over fluorescence intensity maps. Each MCMC sample generated by B-SIM represents a candidate fluorescence intensity map. An appropriate statistical estimator, the mean in this paper, provides the final representation of the fluorescence intensity map, and the extent of sample-to-sample variation provides a measure of confidence in that map.

In Fig. 4a-d, we show reconstructions performed on high-SNR simulated line pairs and experimental HeLa cell images of the previous sections along with line cuts of mean fluorescence intensity map and 50% confidence intervals computed from posterior probability distributions shown as heatmaps. We confirm the existence of features that appear in B-SIM reconstructions in the previous sections based on very low absolute uncertainties where intensity dips occur between line pairs and in the center of loop-like structures in the mitochondria network. In the Extended Data Fig. 1, we show the posterior distributions with confidence intervals for the low-SNR reconstructions of the previous sections, as well as how to use MCMC samples collected by B-SIM to generate a visual representation of a posterior probability distribution.

**Fig. 4:**
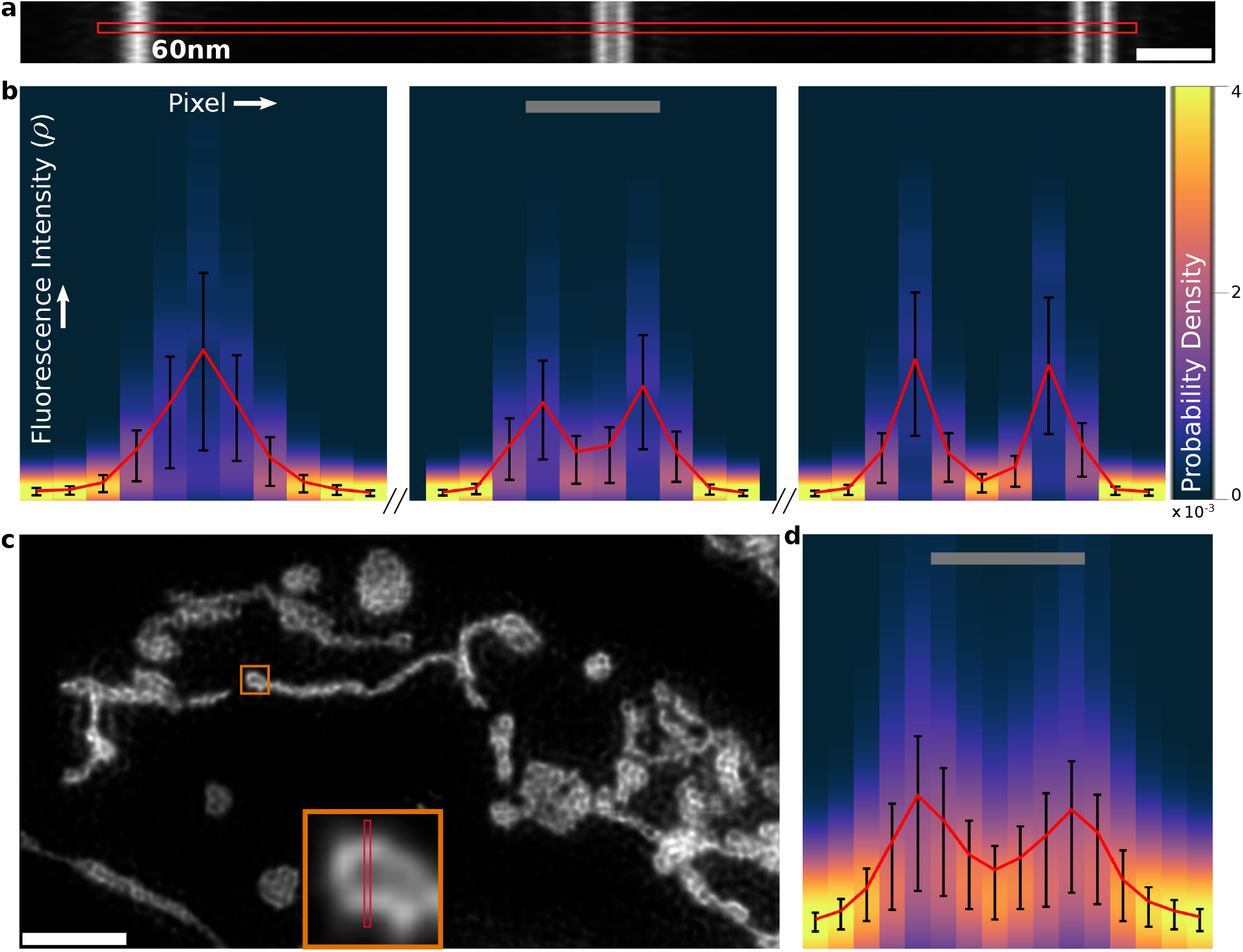
Uncertainty estimation by B-SIM. **a**. B-SIM reconstruction for the high-SNR simulated line pairs of Sec. 2.1. Scale bar 300 nm. **b**. Marginal posterior probability distributions for fluorescence intensity at each pixel along the line cut in a. Mean shown in red along with 50% confidence intervals in black. Scale bar 120 nm. **c**. B-SIM reconstruction of mitochondria networks with high photon counts from Sec. 2.3. Scale bar 2.0 µm. Region of interest (ROI) in the orange box is shown in the inset. **d**. Marginal posterior distributions along the line cut in c. Same color map as in b. Scale bar 195 nm.

In the B-SIM reconstructions presented in this paper, our preference for mean as the estimator is motivated by computational convenience. We find it easier to parallelize the process of generating MCMC samples by placing no expectation that fluorescence intensity maps be smooth, *a priori*. However, due to the ill-posedness arising from diffraction and noise, lack of smoothness constraints results in most MCMC samples predicting pointillistic fluorescence intensity maps as they far outnumber probable smooth maps. With such large degeneracy in the posterior distribution, recovering the most probable sample representing the true (smooth) fluorescence intensity map is a computationally prohibitive task. An efficient alternative is to use an integrative statistical estimator like the mean to recover pixel-to-pixel correlations in biological features warranted by the collected data. Supplementary Fig. S2 shows how the smoothness of the fluorescence intensity map for high-SNR simulated line pairs of Sec. 2.1 improves as the number of MCMC samples used to compute mean is increased.

Additionally, in Fig. S2, we show maps of the inverse of the posterior coefficient of variation (CV), computed as the posterior mean to standard deviation ratio. In the same spirit as SNR for the Poisson distribution, inverse CV can be used to compare reconstruction quality in different regions of the image, and is indicative of the effective SNR we discussed in the introduction resulting from diffraction or the OTF attenuating high frequency information, making SIM reconstruction increasingly ill-posed. Indeed, in the case of simulated line pairs, the inverse CV decreases with separation as shown in Fig. S2. This increase in uncertainty in fluorescence intensity maps results from the increased model uncertainty where numerous candidate fluorescence intensity maps predict raw images nearly identical to the ones produced by the ground truth with similar probabilities. In fact, out of the approximately 400 samples or candidate fluorescence intensity maps collected by B-SIM for the line pairs, the similarity in morphology to the ground truth decreases with separation. While over 50% of the samples have two peaks for the 120 nm separated line pair, a large number of samples predict a merging of the the two lines for the unresolvable 60 nm separated line pair and only about 15% of the samples have two peaks.

## 3 Discussion

We have developed a physically accurate framework, B-SIM, to recover super-resolved fluorescence intensity maps with maximum recoverable resolution from SIM data within the Bayesian paradigm. We achieved this by incorporating the physics of photon shot noise, and camera noise into our model facilitating statistically accurate estimation of the underlying fluorescence intensity map at both high- and low-SNR. Our method standardizes the image processing workflow and eliminates the need for choosing different tools or *ad hoc* tuning reconstruction hyperparameters for different SNR regimes or fluorescence intensity maps. We benchmarked B-SIM on both simulated and experimental data, and demonstrated improvement in contrast and feature recovery in both high- and low-SNR regimes at up to ≈25% shorter length scales compared with conventional methods. We found that B-SIM continues to recover superresolved fluorescence intensity maps with high fidelity at low-SNR, where Fourier methods are dominated by artifacts. Furthermore, because our Bayesian approach recovers a probability distribution rather than a single fluorescence intensity map, we used it to provide absolute uncertainty estimates on the recovered fluorescence intensity maps in the form of posterior variance and inverse CV to compare the quality of reconstruction in different regions. Lastly, we found B-SIM to be robust against phase errors in pattern estimates.

The generality afforded by our method comes at a computational cost that we mitigate through parallelization. If implemented naively, image reconstruction requires the computation of convolutions of the fluorescence intensity map with the PSF every time we sample the fluorescence intensity map from our posterior. This computation scales as the number of pixels cubed. However, the parallelization of the sampling scheme, as we have devised here based on the finite PSF size, reduces real time cost by a factor of the number of cores being used itself, moderated by hardware-dependent parallelization overhead expense.

Moving forward, we can extend and further improve our method in a number of ways. One improvement would be a GPU implementation where parallelization over hundreds of cores may significantly reduce computation time. Furthermore, more efficient MCMC sampling schemes, such as Hamiltonian Monte Carlo [57, 58], and more informative prior distributions over fluorescence intensity maps may be formulated to reduce the number of MCMC iterations needed to generate large number of uncorrelated MCMC samples, accelerating convergence to a high-fidelity mean fluorescence intensity map.

Additionally, the sampling method developed here is generalizable and applicable to significantly improve other computational imaging techniques where the likelihood calculation is dominated by convolution with a PSF. As this describes most microscopy techniques, we anticipate our approach will be broadly useful to the imaging community, particularly dealing with modalities that are often SNR-limited, such as Raman imaging [59, 60]. Alternatively, our approach could be applied to fully-principled image deconvolution and restoration.

In the context of SIM, our method’s extensions to other common experimental settings are also possible. For example, other camera architectures, such as EMCCD, requiring alternative noise models [19] are easily incorporated into our framework. B-SIM’s versatility is also extendable by incorporating a more realistic PSF obtained by calibration measurements, learned from the samples directly, or simulated using a vectorial model to account for refractive index mismatch. One straight forward extension is 3D-SIM, where a straightforward modification of B-SIM to include the more complex 3D-SIM patterns and 3D PSF allow for 3D-SIM reconstruction as shown in Sec. 5.5 and supplementary Fig. S6.

Due to its generality and superior performance at both high- and low-SNR, we expect B-SIM will be widely adopted as a tool for high-quality SIM reconstruction. Furthermore, in the low-noise regime, fully unsupervised B-SIM is competitive with deep learning approaches. However, due to the use of a physics-based model, B-SIM is applicable to arbitrary fluorescent sample structures with no need to tune model parameters or retrain when switching from one class of samples to another, for instance, from mitochondria networks to microtubules.

## 4 Data Availability

Simulation data, experimental data, and scripts used to perform the mcSIM Wiener and FISTA-SIM reconstructions are available online [61]. Wiener and FISTA-SIM reconstruction code is available online, and the version used in this work is archived on Zenodo [62]. B-SIM code is available online [63].

## 5 Methods

Here we first setup an image formation model to generate noisy diffraction limited raw images for SIM where a sample is illuminated using multiple patterns, and then design an inverse strategy to estimate the fluorescence intensity map *ρ*(***r***) in the sample plane. In other words, our goal is to learn the probability distribution over fluorescence intensity maps given the collected raw images and pre-calibrated illumination patterns.

We use Bayesian inference where a probability distribution over fluorescence intensity maps based on predefined domain called prior is updated through a likelihood function that incorporates the experimental data/images.

### 5.1 Image Formation Model

Let *ρ*(***r***) be the fluorescence intensity at each point of space in the sample under a microscope. In SIM, we illuminate the sample a total of L times, with running index l, with sinusoidal spatial patterns given by

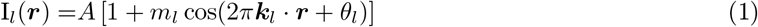

where A is the amplitude, m_*l*_ is the modulation depth, ***k***_*l*_ is the frequency, and θ_*l*_ is the phase of the lth illumination pattern. This illumination causes the sample at each point in space to fluoresce with brightness proportional to its fluorescence intensity multiplied by the illumination at that point. The light from the sample passes through the microscope, and this process is modelled as convolution with the point spread function (PSF) of the microscope, which in this work, we assume to be a 2D Gaussian given by

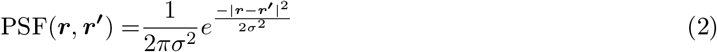

where 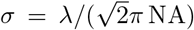 [64], λ is the emission wavelength, and NA is the numerical aperture of the objective lens. A more general PSF is also easily incorporated into our current framework.

The mean number of photons detected by the camera for the lth illumination pattern, ***µ***^*l*^, is the integral of the irradiance over the area of the n-th pixel, A_*n*_,

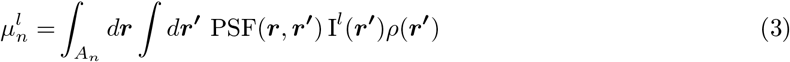

which we write more compactly as

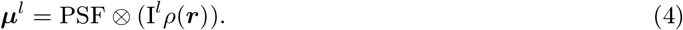

Here ***µ***^*l*^ is the collection of expected brightness values on the camera for the l-th illumination pattern and ⊗ is the convolution operation.

The number of photons detected on each pixel is Poisson distributed

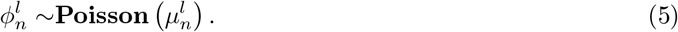

Finally, the camera electronics read out the pixel value and convert the measurement to analog-to-digital units (ADU). The number of ADU are related to the photon number by a gain factor, G_*n*_, and an offset o_*n*_. We model the effect of readout noise as zero mean Gaussian noise with variance 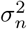. The final readout of the nth pixel, 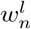, is thus

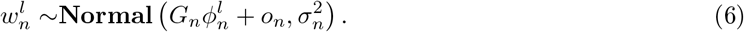

With this observation model for each pixel, we can write the likelihood for a set of observations (raw images) as the product over probabilities of individual raw images given the fluorescence intensity map and camera parameters as

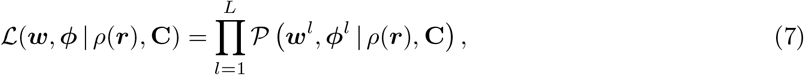

where ***w*** is the collection of readout values on the camera, ***ϕ*** is the collection of photon counts detected by the camera, **C** is the collection of all the camera parameters including the gain factor, offset, and readout noise variance for each pixel, and the vertical bar “|” denotes dependency on variables appearing on the bar’s right hand side.

Now, since the number of photons detected on the camera ***ϕ*** are typically unknown, we must marginalize (sum) over these random variables yielding the following likelihood

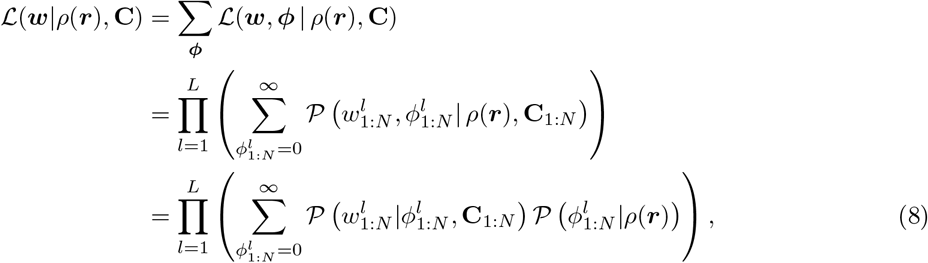

where, in the third line, we have used the chain rule for probabilities and ignored any dependency of photon detections on the camera parameters in the second term. Next, we note that expected values ***µ***^*l*^ for Poisson distributed photon detections are deterministically given by the convolution in equation (4). This constraint allows us to write down the final likelihood expression as

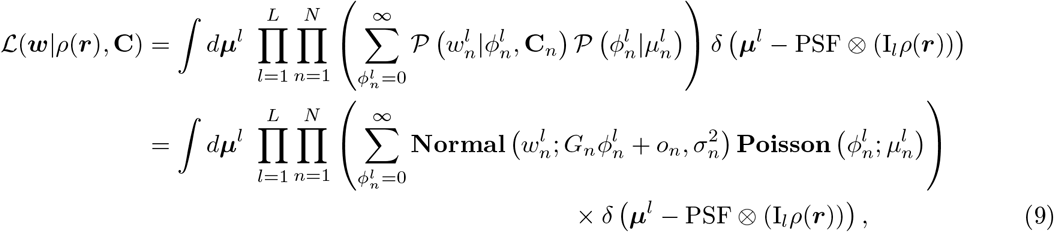

where, assuming each camera pixel to be stochastically independent, we multiply the individual probabilities for camera readout on each pixel. We also note that we haven’t yet chosen a grid in the sample plane on which to discretize the fluorescence intensity map *ρ*(***r***). Such a grid is chosen according to convenience due to the simple additive property of the Poisson distributions that dictate the photons entering a camera pixel. For demonstration purposes, we assume that the fluorescence intensity map is Nyquist sampled on a grid twice as fine as the camera pixel grid. We denote this fluorescence intensity map on the m-th point on this grid with *ρ*_*m*_ and the collection of these values with ***ρ***.

Now, with the likelihood at hand, we move on to the formulation of an inverse strategy to estimate the fluorescence intensity map ***ρ***.

### 5.2 Inverse Strategy

Using Bayes’ theorem, we now construct the posterior distribution over fluorescence intensity maps 𝒫 (***ρ*** | ***w*, C**) from the product of the likelihood function and a suitably chosen prior probability distribution. That is,

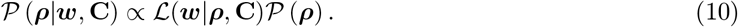

We have already described the likelihood in the last subsection.

In order to select priors, we note that as more observations (images) are incorporated into the likelihood, the likelihood dominates over the prior. In effect, Bayesian inference updates the prior through the likelihood yielding a posterior. Therefore, priors and posterior distributions should have the same support (parameter space domain). A convenient prior for ***ρ*** is an uncorrelated product of Gamma distributions

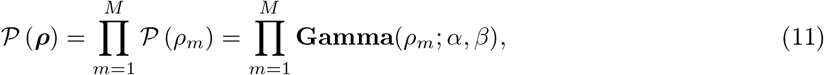

where we choose the shape parameter α = 0.1 and scale parameter β = 10 so that the variance is large, thereby reducing the influence of prior [65]. Taken together, our full posterior becomes

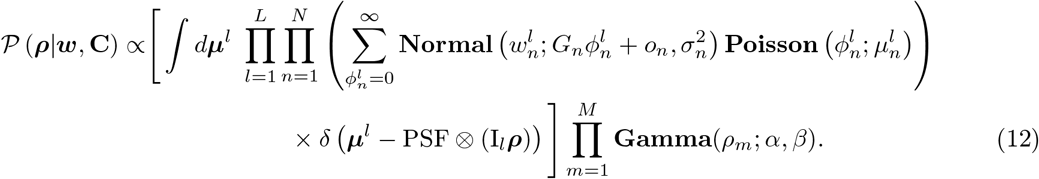

This posterior does not have an analytical form amenable to direct sampling. Therefore, we employ iterative Monte Carlo techniques such as Markov Chain Monte Carlo (MCMC) to generate samples from this posterior. We now describe our MCMC strategy below.

### 5.3 Sampling Strategy: Parallelization

Naively, one may employ the most basic MCMC technique where samples for fluorescence intensity at each pixel *ρ*_*m*_ are generated sequentially and separately using a Gibbs algorithm. To do so, we typically first expand the posterior of equation (12) using the chain rule as

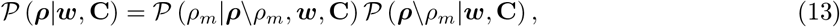

where the backslash after ***ρ*** indicates exclusion of the immediately following parameter *ρ*_*m*_. In this last equation, the first term on the right is the conditional posterior for *ρ*_*m*_ and the second term is considered a proportionality constant for the Gibbs step as it is independent of *ρ*_*m*_. Similarly, we decompose the prior in equation (11) 𝒫 (***ρ***) into a constant 𝒫 (***ρ***\*ρ*_*m*_) and a function of the random variable of interest 𝒫 (*ρ*_*m*_). Plugging these decompositions into equation (12), we arrive at the conditional posterior for *ρ*_*m*_ as

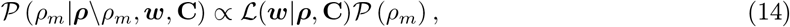

where the first term on the right-hand side is the likelihood of equation (9) as before. Plugging in the expression for the likelihood function into this equation, we get

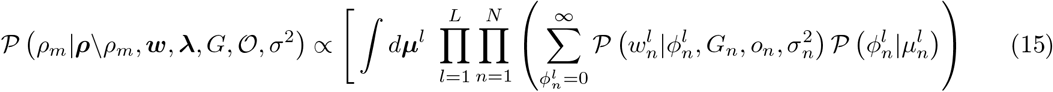

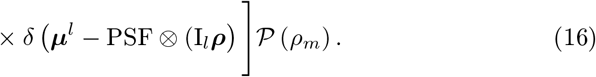

Now, since direct summation over the unobserved photon emissions 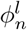 in the likelihood above is intractable, we again may use Monte Carlo techniques and simulate the probabilistic effect of this summation by sampling these intermediate (latent) variables 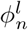. In other words, we further decompose the conditional posterior in the Gibbs step of equation (16) into two steps:

1) sample 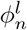 from its conditional posterior

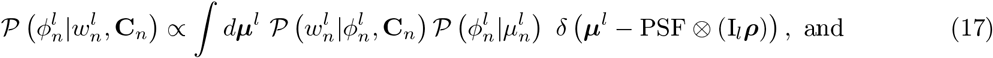

2) sample *ρ*_*m*_ from its conditional posterior

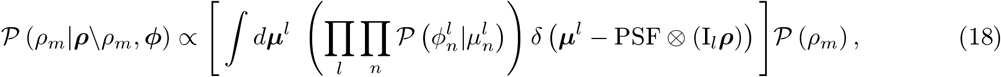

where terms involving ***w*** have disappeared as they only depend on ***ϕ***, which is a fixed quantity in this step. Since these conditional posteriors are again not amenable to direct sampling, we may employ Metropolis-Hastings to accept or reject randomly proposed samples. For instance, a new sample for fluorescence intensity 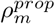 may be proposed from a proposal distribution 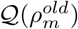 and accepted based on the acceptance probability

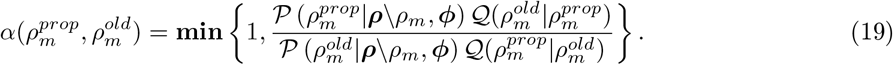

Based on this sampling strategy above, a chain of MCMC samples, initialized with a randomly generated fluorescence intensity map, can be generated. However, this strategy is prohibitively expensive for large images. For instance, sampling fluorescence intensity map defined on a 2048 × 2048 pixel grid once would require computing convolution integrals of equation (4) approximately 4 million times.

A more reasonable approach follows by first realizing that pixels far apart are uncorrelated when the PSF is only a few pixels wide. This allows us to reasonably assume that the fluorescence intensity at a pixel only contributes light in its neighborhood. Consequently, the likelihood ratios in the Metropolis-Hastings step are now approximated using ratios of local likelihoods. This procedure reduces the computational cost by allowing parallelization of the sampling method. More formally,

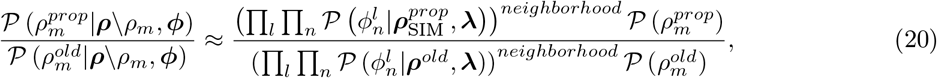

where the superscript “neighborhood” indicates that the likelihood is computed only using pixels in the neighborhood of the fluorescence intensity *ρ*_*m*_. This procedure now allows us to replace the convolution of a 2048×2048 fluorescence intensity map with a much smaller integral performed on a small grid determined by the size of the PSF, typically of order 24 × 24 for Nyquist sampled images. When implementing this strategy on a computer, an appropriate padding, zeros in this paper, must be added as well beyond the boundaries of the image and the parallelized chunks to facilitate computation of convolution integrals for boundary pixels. To validate this strategy, in supplementary Fig. S3, we show posteriors for five MCMC chains with increasing parallelization, generated by B-SIM for the same set of raw images to demonstrate no significant deviations in posteriors for the parallelized versions from the posterior for the non-parallelized version.

Finally, it is well-known that MCMC samples remain correlated for a significant number of iterations [65, 66]. Therefore, to efficiently collect uncorrelated samples, we repeatedly apply simulated annealing [67] where we artificially widen the shape of the posterior distribution at regular intervals and perturb the MCMC chain of samples by accepting improbable samples, as shown in supplementary Fig. S4. The samples at the end of each annealing cycle are then collected together for further analysis.

### 5.4 Computational Expense

For all the examples presented here, we used a 64-core computer where B-SIM takes about a day to compute a 800 × 800 pixel super-resolved fluorescence intensity map, for which we collect around 400 uncorrelated Monte Carlo samples to compute the mean. Furthermore, our method has low memory requirements as only the fluorescence intensity map to be learned and raw images are kept in memory.

### 5.5 3D-SIM

The inference strategy presented above for 2D-SIM largely applies directly to 3D-SIM though the convolution operation in the image formation model needs modification to allow for generation of a stack of images along the microscope’s axial direction [68]. Briefly, we consider a fluorescent sample moved to different locations along the axial direction for patterned illumination, generated by three-beam interference, fixed with respect to the microscope objective [68]. Consequently, in the convolution appearing in the image formation model, the illumination pattern now cannot naively be multiplied with the fluorescence intensity *ρ* in a point-by-point manner as was previously done, *c*.*f*., equation 4. Instead, the axial component of the illumination pattern now multiplies the PSF, as follows

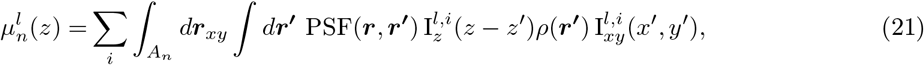

where x and y are the coordinates in the camera plane as before, however, z represents the axial location of the sample. Furthermore, we have assumed that each illumination pattern can be written as a sum of separable axial and lateral sinusoidal components, that is, 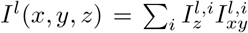 [68]. Fifteen such illumination patterns are generated by using three-beam interference at 3 different angles and 5 phase shifts [68]. Finally, we can write the equation above more compactly as

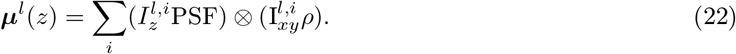

As a proof of principle, we show 3D-SIM reconstruction of simulated spherical shells in supplementary Fig. S6.

### 5.6 Calibrating Camera Parameters

We precalibrate the illumination and camera parameters in equation (12). To determine the read noise, offset, and gain we collect uniform illumination profiles at several different intensity levels and apply the approach of [69].

### 5.7 Decorrelation Analysis

We performed decorrelation analysis using the imageJ plugin provided in [56]. We used default settings: sampling of decorrelation curve N_*r*_ = 50, number of high-pass images considered N_*g*_ = 10, radius min = 0, radius max = 1.

### 5.8 HiFi-SIM

HiFi-SIM reconstructions were performed using the Matlab GUI [29] using default attenuation strength of 0.9 in the “non-Pro” mode.

For the low-SNR ArgoSIM data, HiFi-SIM did not accurately estimate the illumination pattern parameters, most likely due to the highly-structured Fourier domain structure of the line pairs image.

### 5.9 Sample Preparation

#### 5.9.1 Simulated line pairs

Simulated line pairs separated by 0 nm to 390 nm were generated on a grid with pixel size 30 nm. Line pairs were separated by 2.1 µm and each individual line had a flat-top fluorescence intensity profile with width of one pixel. For convenience, line pairs were generated parallel to the pixel grid. We assumed sinusoidal SIM patterns rotated at angles *θ* of 31.235°, 91.235°, and 121.235°. At each angle, the SIM phases were 0°, 120°, and 240°. The SIM frequencies were set at 90 % of the detection band-pass frequency (f_*x*_, f_*y*_) = (cos θ, sin θ) × 0.9 × λ/2 NA. Here we take λ = 500 nm and NA = 1.49.

To generate the simulated data, we multiplied the ground truth image with the SIM patterns and a scaling factor to give the desired final photon number, then convolved the result with a PSF generated from a vectorial model using custom Python tools [70, 71]. The pixel values of this image represent the mean number of photon counts that would be collected on that pixel. To generate a noisy image, we draw from an appropriate Poisson distribution on each pixel. Next we apply camera gain of 0.5 e^*−*^/ADU, offset of 100ADU, and Gaussian read-out noise of standard deviation 4ADU. Finally, we quantize the image by taking the nearest integer value. We generated simulated images at a range of maximum photon numbers ranging from 1 photons to 10 000 photons.

In the main text, we display datasets with nominal photon numbers 10 000 and 300. We performed Wiener SIM reconstruction using the Wiener parameter w = 0.3 [18]. For the FISTA-SIM, the total variation regularization strength was 1 × 10^*−*6^ and 1.7 × 10^*−*7^ [48].

#### 5.9.2 Argo-SIM calibration slide

Argo-SIMv1 (Argolight) slide SIM images were acquired on a custom Structured Illumination Microscope [18] using a 100x 1.3 NA oil immersion (Olympus, UPlanFluor) objective and 465 nm excitation light derived from a diode laser (Lasever, OEM 470 nm-2 W). The effective pixel size was 0.065 µm. The SIM patterns are generated using a digital micromirror device (DMD) in a conjugate imaging plane. The DMD does not fill the camera field of view. The illumination profile is nominally flat because the DMD is illuminated by the excitation light after it passes through a square-core fiber. Laser speckle is suppressed by rapidly shaking the optical fiber using a fiber shaker based on the design of [72]. Images were acquired on an Orca Flash4.0 v2 (C11440-22CU) Hamamatsu sCMOS camera with gain of ≈0.51 e^*−*^/ADU, offset of 100 ADU, and RMS readout noise of 2 e^*−*^. The camera exposure time was fixed at 100 ms, and the signal level was varied by changing the illumination time using the DMD as a fast shutter. The illumination times used were 100 ms, 30 ms, 10 ms, 3 ms, 1 ms, and 0.3 ms.

In the main text the high-SNR dataset used 100 ms illumination time, while the low-SNR dataset used 3 ms. We performed Wiener SIM reconstruction using w = 0.1 and 0.2 respectively. We performed FISTA-SIM with TV strength 1 × 10^*−*7^ and 1.7 × 10^*−*8^.

The “gradually spaced line” test patterns were selected from the many available patterns on the Argo-SIM slide. These patterns consist of 14 line pairs with spacings of 0 nm to 390 nm in 30 nm steps which are arranged in a ≈36 µm × 36 µm square.

The SIM patterns are generated using lattice vectors 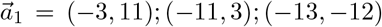 and 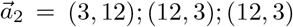 respectively. This results in SIM patterns at ≈71 % of the excitation pupil [18]. Once the Stokes shift is accounted for, this allows for resolution enhancement by a factor of ≈1.8.

#### 5.9.3 Live HeLa cells with labelled mitochondria

HeLa cells (Kyto strain) were grown in 60 mm glass petri dishes on Poly-d-lysine coated 40 mm #1.5 coverslips (Bioptechs, 40-1313-03192) for a minimum of 48 h in DMEM media (ATCC 30-2002) supplemented with 10 % FBS (ATCC 30-2020) and 1 % Penicillin-Streptomycin solution (ATCC, 30-2300) at 37 °C and 5 % CO2. Cells were live stained with 200 nM Mitotracker Deep Red (ThermoFisher, M22426) in DMEM medium for 15 min in the same incubation environment. The staining solution was then aspirated off and fresh DMEM medium was added for 5 min to rinse. The sample coverslip was then transferred to an open-top Bioptechs FCS2 chamber and imaged in pre-warmed (37 °C) DMEM culture medium.

HeLa cells with labelled mitochondria were imaged on the same instrument described for the Argo-SIM calibration slide using 635 nm excitation light derived from a diode laser (Lasever, LSR635-500). The camera integration time was fixed at 100 ms and the signal level was varied by changing the illumination time using the DMD as a fast shutter. Illumination times were 100 ms, 10 ms, 1 ms, and 0.2 ms.

In the main text the high-SNR dataset used 100 ms illumination time, while the low-SNR dataset used 10 ms. We performed Wiener SIM reconstruction using w = 0.1 and 0.3 respectively. We performed FISTA-SIM with TV strength 1 × 10^*−*7^ and 5.5 × 10^*−*8^.

The SIM patterns are generated using lattice vectors 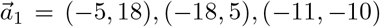, and 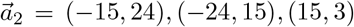 respectively. This results in SIM patterns at ≈71 % of the excitation pupil [18]. Once the Stokes shift is accounted for, this allows for resolution enhancement by a factor of ≈1.8.

#### 5.9.4 Microtubules

SUM159 were maintained in a tissue culture incubator at 37°C with 5% CO_2_. The culturing media comprised F-12/Glutamax (Thermo Fisher Scientific) medium supplemented with 5% fetal bovine serum (Gibco), 100 U/mL penicillin and streptomycin (Thermo Fisher Scientific), 1 µg mL^*−*1^hydrocortisone (H-4001; Sigma-Aldrich), 5 µg mL^*−*1^ insulin (Cell Applications), and 10 mM 4-(2-hydroxyethyl)-1-piperazine-ethane-sulfonic acid (HEPES).

For imaging microtubules, SUM159 cells were seeded onto a 35mm #1.5 glass bottom dish (Mat-Tek Life Sciences) and, once reached 60% confluency, treated with SPY555-tubulin (Cytoskeleton Inc, Cat: Cy-SC203) following the manufacturer’s instructions. Live cell imaging experiments were performed in L15 imaging medium (Thermo Fisher Scientific) supplemented with 5% serum and 100 U/ml penicillin/streptomycin.

Microtubules were imaged using a TIRF-SIM system custom-built on a Nikon Inverted Eclipse TI-E microscope [73]. Briefly, the structured illumination is generated using 488 nm and 561 nm (300 mW, Coherent, SAPPHIRE LP) lasers, an acousto-optic tunable filter (AOTF; AA Quanta Tech, AOTFnC-400.650-TN), an achromatic half-wave plate (HWP; Bolder Vision Optik, BVO AHWP3), a ferroelectric spatial light modulator (SLM; Forth Dimension Displays, QXGA-3DM-STR) and a 100X 1.49NA objective (Olympus UAPON 100XOTIRF). The images are acquired using Prime BSI Express sCMOS camera (Teledyne Photometrics) at a 18 frames/s rate, using 20 ms/frame exposure.

## Author Contributions

SP and DPS conceived of the project. AS, JSB, and ZF carried out mathematical derivations. AS developed all original code. PTB provided datasets for the simulated line pairs, Argo-SIM slide, and HeLa cells and performed FISTA-SIM reconstructions. CT and CK provided microtubule datasets. RK prepared the biological samples. AS and PTB wrote the paper. DPS and SP oversaw all aspects of the project.

## Acknowledgements

We thank Miyeko Mana for the gift of the mitochondrial stain. We thank Ke Hu, John Murray, Mohamadreza Fazel, and Maxwell Schweiger for helpful discussions. PTB and DPS acknowledge funding support from the NIH (RF1MH128867) and Scialog, Research Corporation for Science Advancement, and Frederick Gardner Cottrell Foundation (28041). CK acknowledges support from the NSF Faculty Early Career Development Program (award number: 1751113) and the NIH (R01GM127526). SP acknowledges support from the NIH (R01GM134426, R01GM130745, and R35GM148237).

## Supplementary Information

**Fig. S1:**
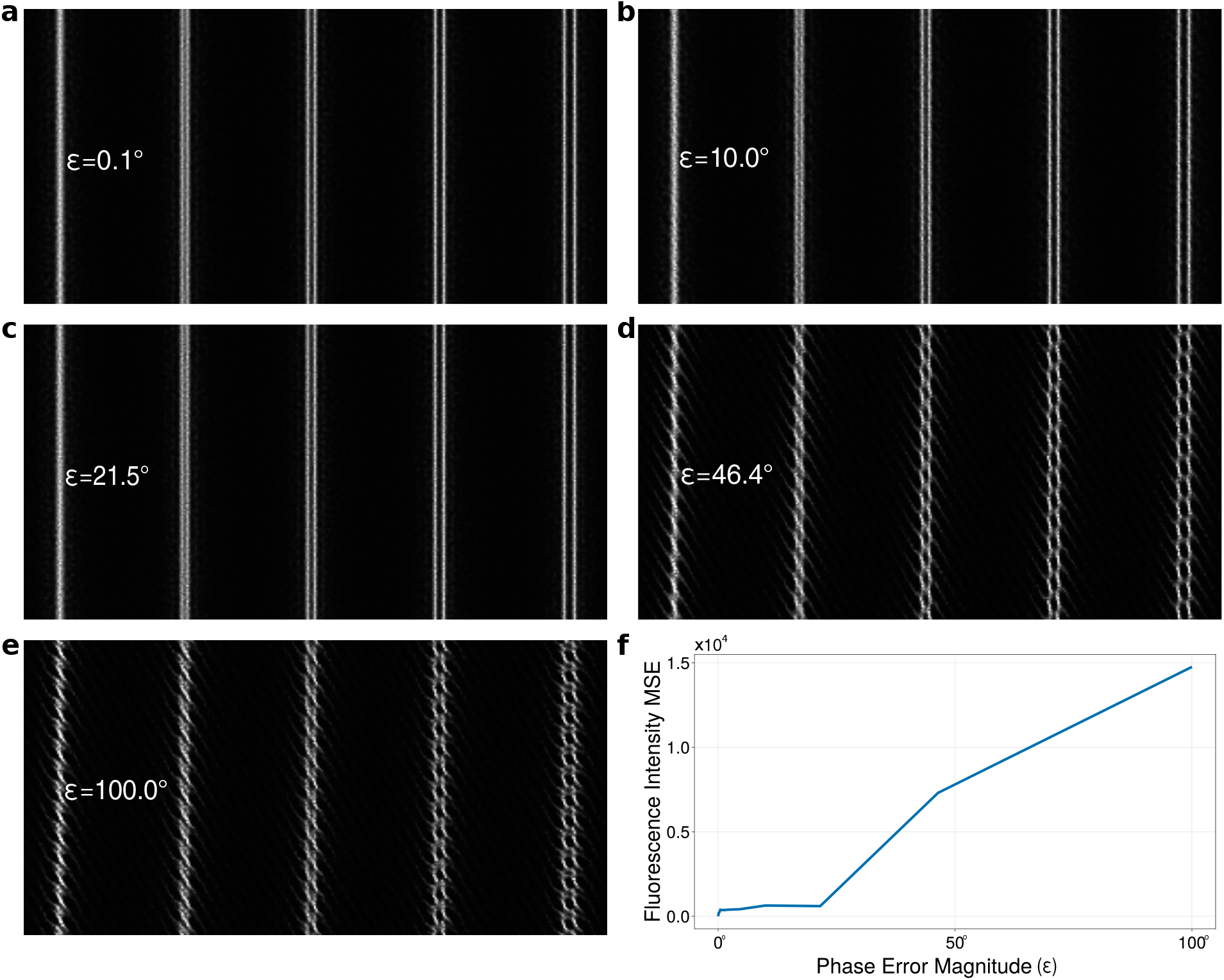
B-SIM reconstruction with increasing illumination pattern phase error ε. **a-e**. B-SIM reconstructions obtained by adding phase errors to patterns for a single orientation. The errors vary in magnitude from 0.1° to approximately 100°. **f**. Mean square error in B-SIM reconstruction. Significant reconstruction artifacts only appear in d when the error is as large as 45°, indicating robustness of B-SIM reconstructions with respect to small errors in pattern estimates.

**Table S1:** Mean-square error (MSE) and peak signal-to-noise ratio (PSNR) for SIM reconstructions of simulated line pairs at both high and low SNR.

|  | high SNR |  | low SNR |  |
| --- | --- | --- | --- | --- |
|  | MSE | PSNR | MSE | PSNR |
| mcSIM | 0.019 | 17.3 | 0.021 | 16.8 |
| HiFi-SIM | 0.016 | 18.1 | 0.017 | 17.8 |
| FISTA-SIM | 0.014 | 18.7 | 0.016 | 17.9 |
| B-SIM | 0.014 | 18.6 | 0.014 | 18.4 |

**Fig. S2:**
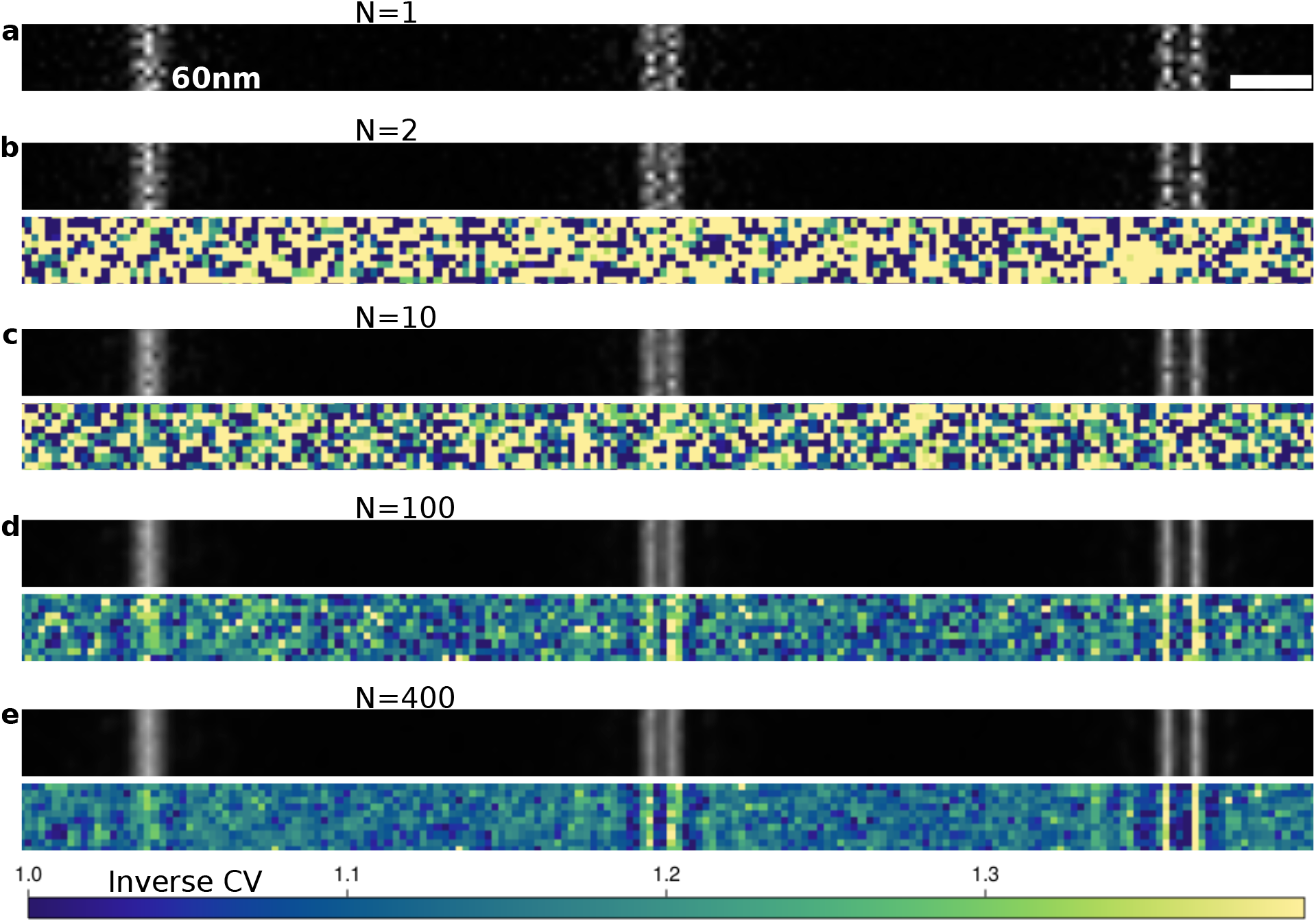
B-SIM reconstruction smoothness as function of number of MCMC samples. **a**. A single MCMC sample generated by B-SIM for the high-SNR simulated line pair of Sec. 2.1. SIM reconstructions of MitoTracker deep red labelled mitochondria in HeLa cells at high-SNR. Scale bar 300 nm. **b-e**. Mean fluorescence profile as the number of samples are increased along with ICV maps.

**Fig. S3:**
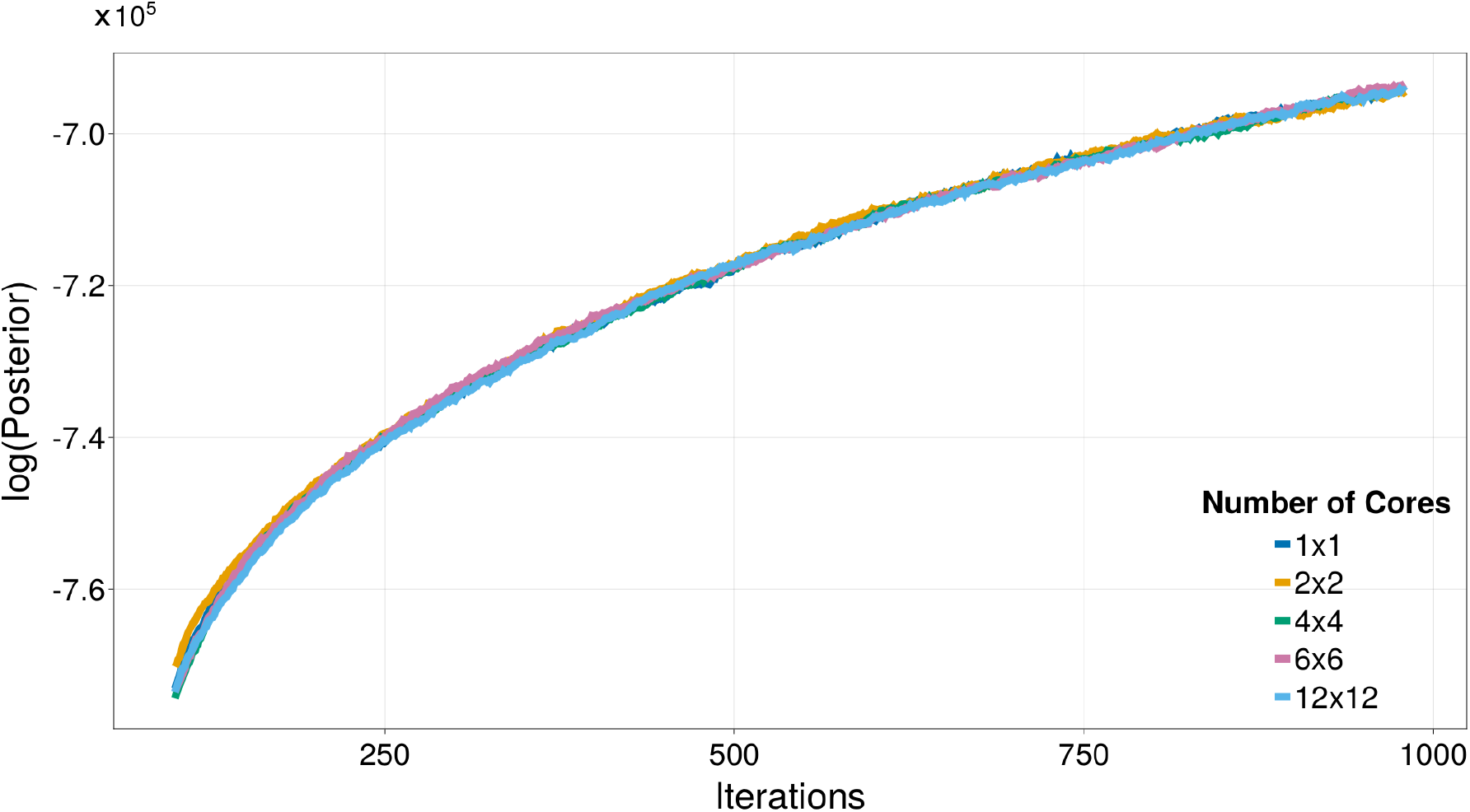
Validity of parallelized MCMC sampling based on local likelihoods. The logarithm of the posterior for the same section of five MCMC chains with increasing parallelization, as generated by B-SIM for an 84 × 84 crop of the high-SNR raw images for the simulated line pairs. All the chains have significant overlap indicating that parallelization based on the finite width of the PSF doesn’t introduce significant errors.

**Fig. S4:**
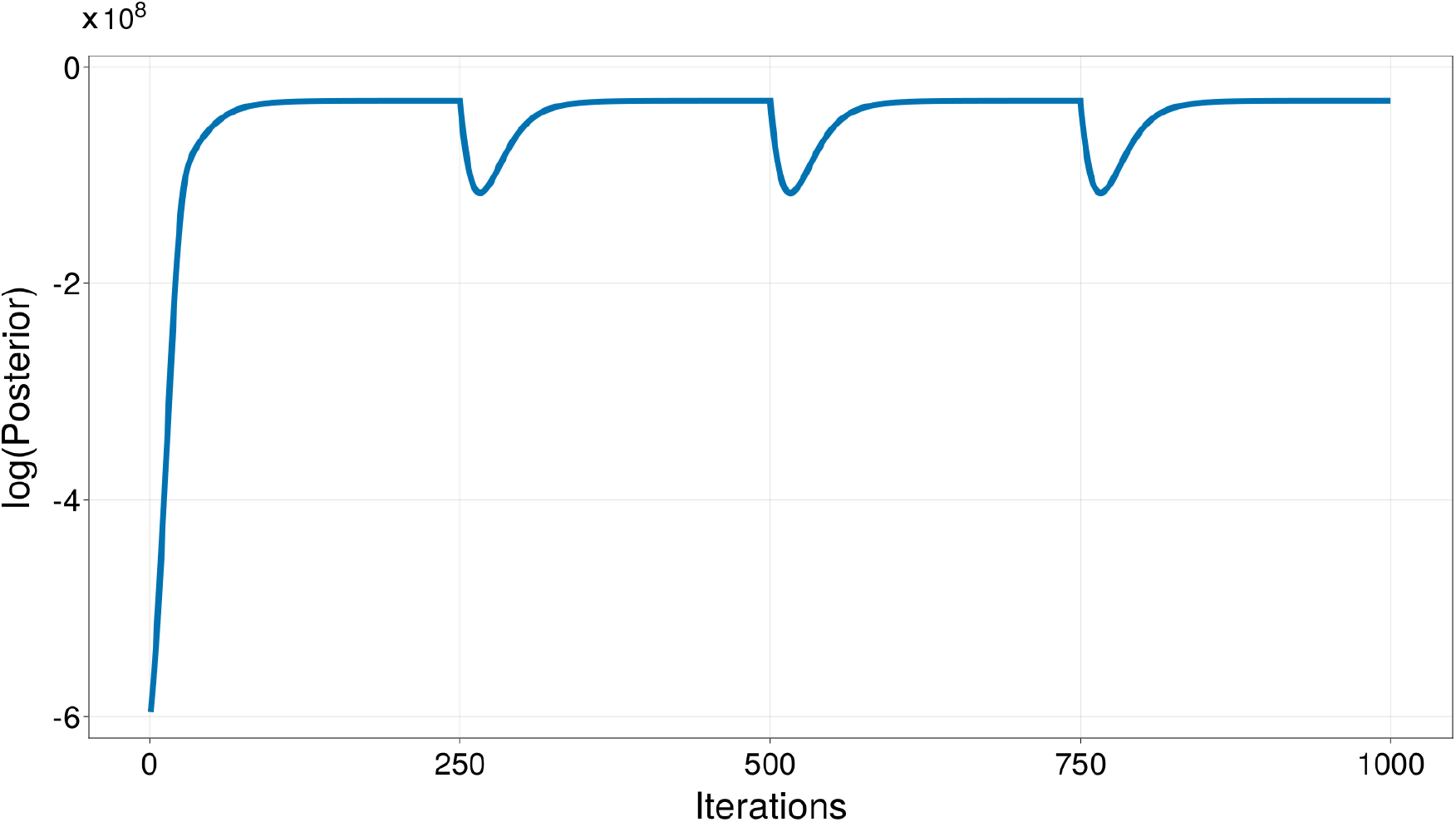
Logarithm of sample-to-sample posterior along a B-SIM MCMC chain. MCMC samples are typically collected only after the initial convergence period, also known as burn-in, which here lasts from iteration 0 to 100 approximately. Most samples during burn-in have very low posterior probabilities and are therefore ignored to avoid any bias. We avoid correlations among samples by artificially widening the posterior using simulated annealing at regular intervals (every 250 iterations here), and let the chain first move away from the converged value and then return to a different sample upon convergence. The samples at the end of each annealing cycle are then used to compute the mean fluorescence intensity map.

**Fig. S5:**
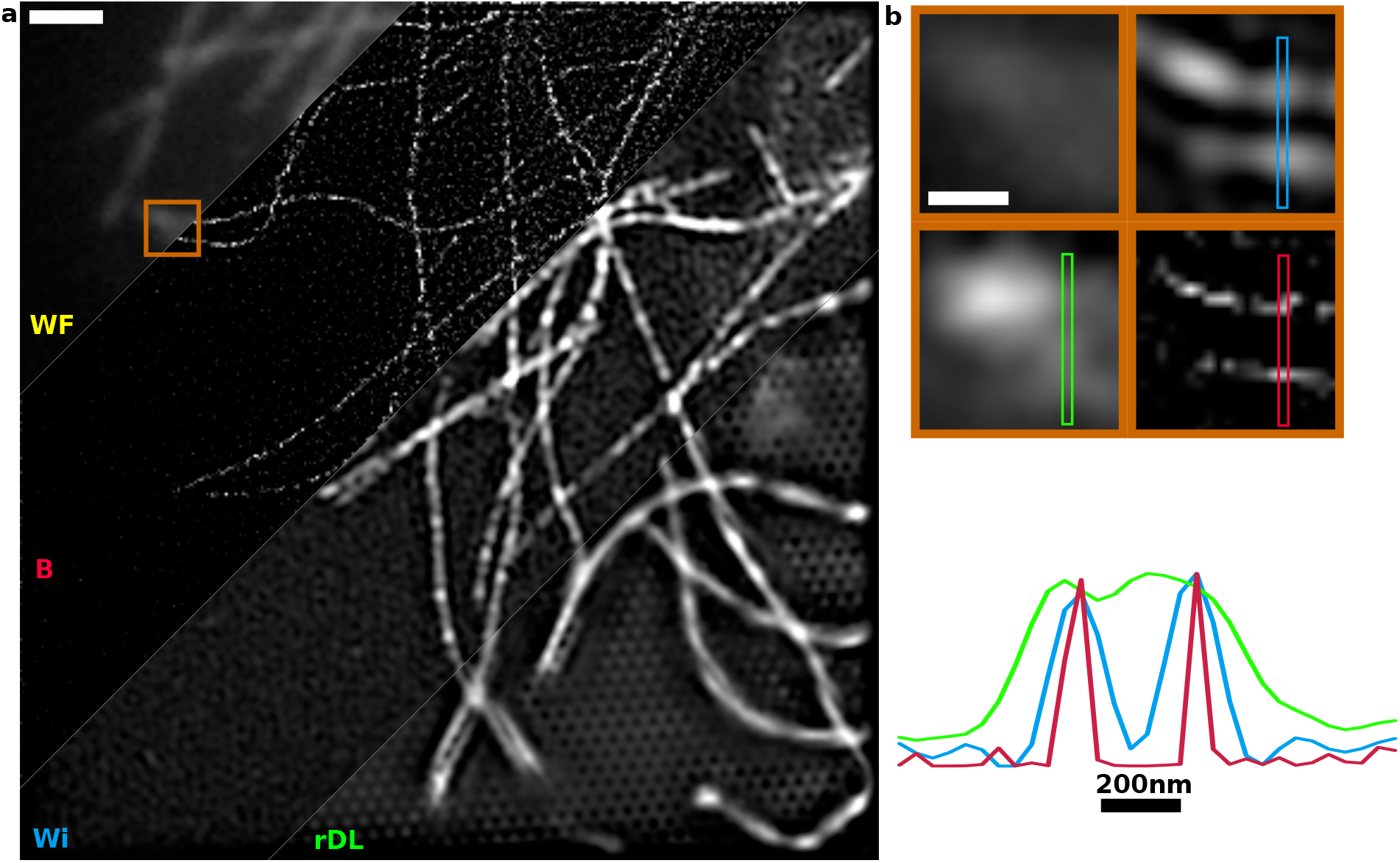
SIM reconstruction of microtubules. **a**. Pseudo-widefield (WF) and SIM reconstructions of SPY555-tubulin labelled microtubules in SUM159 cells. We compare direct reconstruction by Wiener filter (Wi) based HiFi-SIM, deep learning based rDL-SIM denoising followed by HiFi-SIM reconstruction (rDL), and B-SIM (B) reconstruction. Scale bar 1.0 µm. **b**. Region of interest corresponding to the orange box in a. (top) shown in widefield, rDL-SIM, Wiener, and B-SIM. Scale bar 300 nm. Line cuts (bottom) demonstrate that B-SIM provides the highest contrast in reconstructions. Furthermore, while all reconstructions have some noise amplification due to significant out-of-focus background, rDL-SIM reconstruction suffers from significant artifacts including hammerstroke and artificial discontinuities in microtubules.

**Fig. S6:**
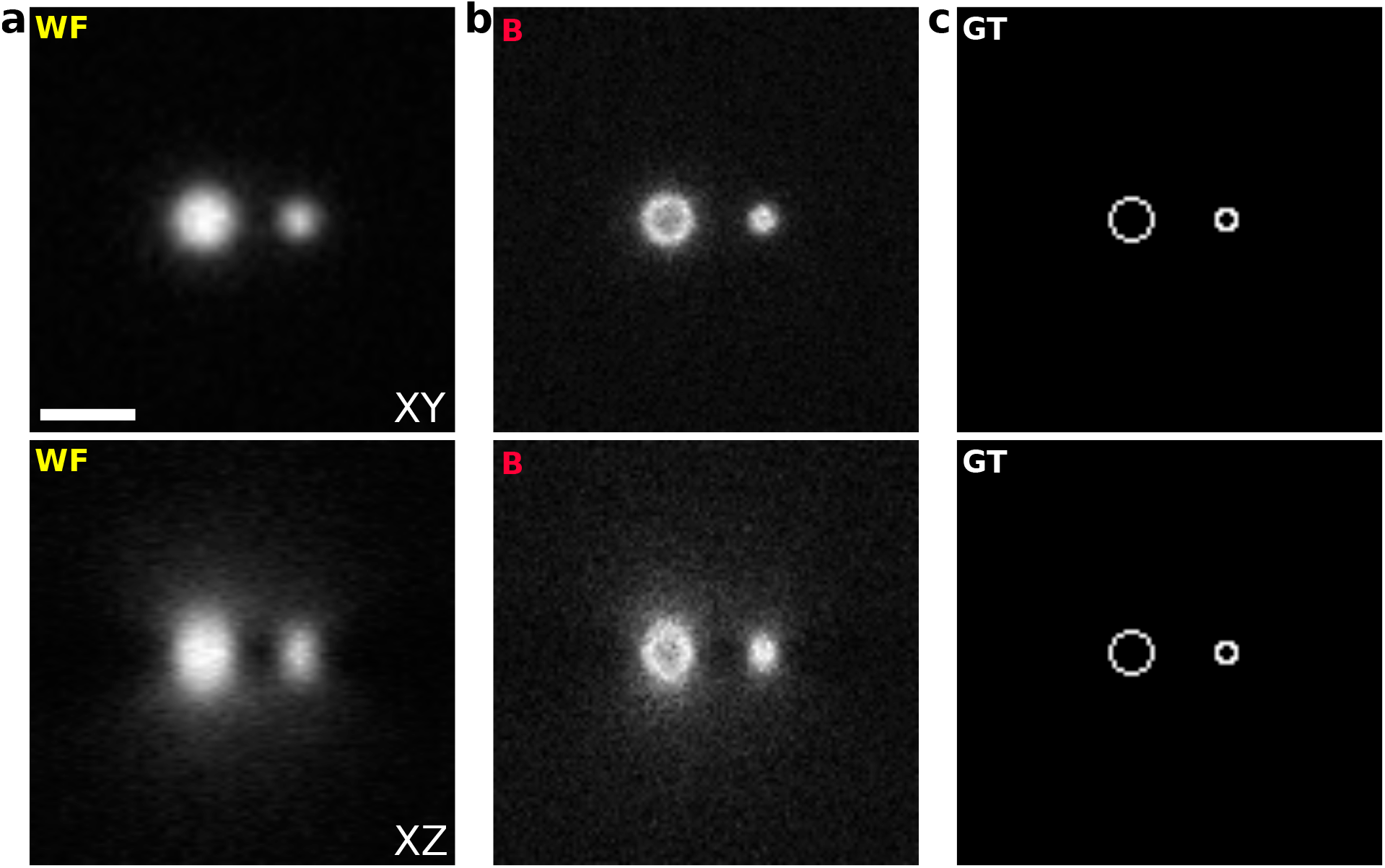
B-SIM reconstruction for simulated 3D-SIM data. The stack of images is generated along the axial direction for an optical setup with NA=1.49 objective, 500nm wavelength light, and axial separation of 30nm between each set of planes. The simulated fluorescent sample contains two 30nm thick shells of diameter 90nm and 210nm. **a**. Pseudo-widefield (WF) images, along an x-y plane (top) and an x-z plane (bottom). **b**. B-SIM reconstruction (B). **c**. Ground Truth (GT). Scale bar 600nm. Here, B-SIM is able to resolve the 90nm sphere along the x-y plane using the Sparrow criterion with a central dip in intensity, while only the 210nm sphere is resolved along the axial direction.

